# Rational construction of compact de novo-designed biliverdin-binding proteins

**DOI:** 10.1101/453779

**Authors:** Molly M. Sheehan, Michael S. Magaraci, Ivan A. Kuznetsov, Joshua A. Mancini, Goutham Kodali, Christopher C. Moser, P. Leslie Dutton, Brian Y. Chow

## Abstract

We report the rational construction of a *de novo*-designed biliverdin-binding protein by first principles of protein design, informed by energy minimization modeling in Rosetta. The self-assembling tetrahelical bundles bind biliverdin IXa (BV) cofactor auto-catalytically *in vitro*, similar to photosensory proteins that bind BV (and related bilins, or linear tetrapyrroles) despite lacking sequence and structural homology to the natural counterparts. Upon identifying a suitable site for cofactor ligation to the protein scaffold, stepwise placement of residues stabilized BV within the hydrophobic core. Rosetta modeling was used in the absence of a high-resolution structure to define the structure-function of the binding pocket. Holoprotein formation indeed stabilized BV, resulting in increased far-red BV fluorescence. By removing segments extraneous to cofactor stabilization or bundle stability, the initial 15-kilodalton *de novo*-designed fluorescence-activating protein (“dFP”) was truncated without altering its optical properties, down to a miniature 10-kilodalton “mini,” in which the protein scaffold extends only a half-heptad repeat beyond the hypothetical position of the bilin D-ring. This work demonstrates how highly compact holoprotein fluorochromes can be rationally constructed using *de novo* protein design technology and natural cofactors.

## INTRODUCTION

*De novo*-designed proteins are powerful tools for exploring the principles and expanding the boundaries of protein folding, protein-protein interaction, and protein biochemical functions that build on structure-function and protein sequence diversity landscapes that are entirely distinct from those of natural protein scaffolds^1-4^. Self-assembling tetrahelical bundles^5-15^, which are created by simple binary patterning of hydrophobic and hydrophilic residues with high α-helical propensity^16^, comprise the best established class of *de novo*-designed scaffolds. These scaffolds can provide stable single-chain frames into which biological cofactors can bind, as designed protein maquettes^6-10^ that are useful for rationally engineering artificial holoproteins because the cofactor-interacting function of an individual residue within the core is largely isolated from the functions of the others (**Figure 1a**).

**Figure 1.**
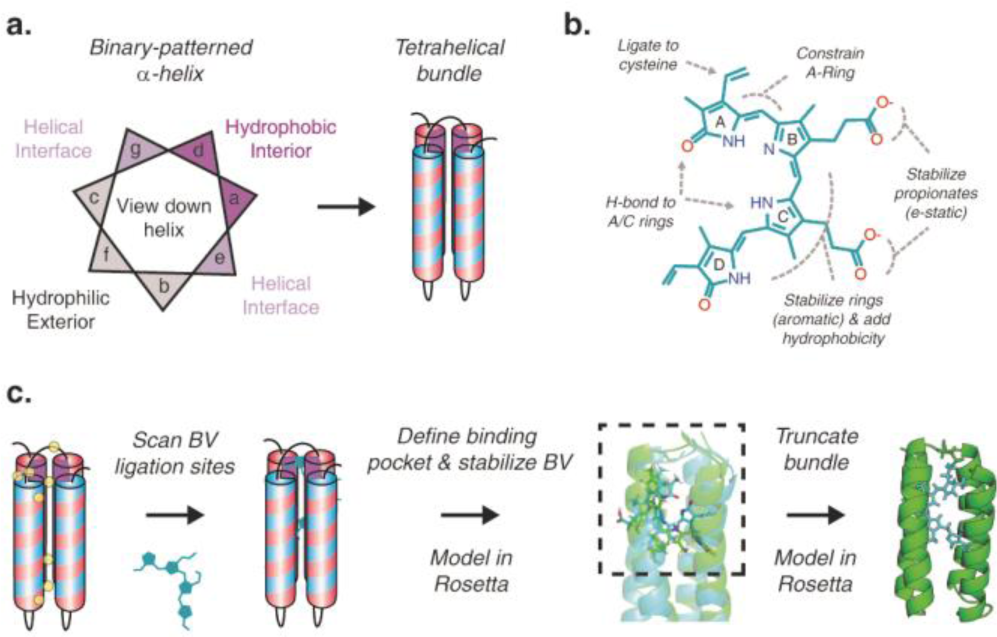
Engineering *de novo*-designed proteins to stabilize biliverdin. **(a)** Self-assembling single-chain tetrahelical bundles created by binary patterning of hydrophobic and hydrophobic residues of high propensity for α-helix formation, as described by the helical wheel. **(b)** Rational strategy for stabilizing biliverdin (BV) within the core. **(c)** Step-wise construction of the *de novo*-designed holoprotein.

Previously reported maquette holoproteins have typically incorporated rigid and planar cofactors, such as hemes, chlorins, porphyrins, and flavins. Recently, we reported that maquettes can also bind highly flexible bilins or linear tetrapyrroles, and identified structure-function determinants for effective autocatalytic ligation of phycocyanobilin (PCB), namely the presence of a free cysteine and the stabilization of the bilin propionates^9^.

Here, we report the rational construction of compact *de novo*-designed proteins that bind biliverdin (BV), the optically active cofactor in natural photosensory bacteriophytochromes (Bph) and Bph-derived protein tools^17-24^. Energy minimization modeling in Rosetta^13-14, 25-27^ informed the strategic placement of residues that stabilized BV within the core and consequently increased its far-red fluorescence, despite lacking sequence or structural homology to natural biological fluorochromes. By eliminating structural vestiges found when engineering from natural scaffolds, bili-proteins as small as 10-kD molecular weight were successfully forward-engineered, which for context is half the size of a minimal fluorescent domain engineered from a Bph^20^.

## RESULTS AND DISCUSSION

### Rational design and construction strategy

Structure-function insights on cofactor stabilization in reported engineered bili-proteins can inform rational approaches for constructing *de novo*-designed ones. To date, fluorescent proteins (FPs) have been evolved from natural bacteriophytochromes (Bph)^17-23^, phytochromes (Phy)^28-30^, allophycocyanin light-harvesting complexes^31-33^ (AP), and fatty acid-binding muscle proteins^34-35^ (FABP). These engineered proteins follow the general principle of rigidifying the protein (i) to stabilize the floppy cofactor in a fluorescent conformation, (ii) to limit solvent and oxygen access to the cofactor, and (iii) to prevent the intrinsic protein structural re-arrangements associated with their natural signaling roles.

Specific insights from reported crystal structures of Bph‐ and Phy-derived fluorescent proteins^22-23, 28^ led to a maquette design strategy for stabilizing the bilin by hydrogen bonding to the propionates and A-ring of the linear tetrapyrrole, plus the addition of hydrophobic core bulk around the cofactor D-ring (**Figure 1b**). Accordingly, in our rational construction strategy (**Figure 1c**), we first experimentally identified a suitable cysteine site for cofactor attachment to an initial scaffold derived from a family of maquettes with molten globular cores^7^,^36^ that accommodate a range of cofactor types and sizes^37^. Cofactor-stabilizing residues were subsequently introduced in a step-wise manner to define a pocket within the apo bundle core. Binding pocket structure-function was informed by energy minimization modeling using Rosetta, which was employed given the absence of a high-resolution structure for this scaffold and the demonstrated ability of the approach for predicting tetrahelical bundle structures and their binding sites to a rigid/planar cofactor^13-15^.

### Cysteine ligation scanning

Bilins can be covalently attached *in vitro* to reconstitute holoproteins because their ligation to cysteine residues is autocatalytic ^9, 38-39^. To systematically identify suitable ligation positions around which to construct a binding pocket, we first scanned cysteine sites throughout the core and loops for BV covalent attachment efficiency to bacterially produced proteins *in vitro* (**Figure 2a-b**). In these cysteine-scan experiments, all core residues (a and d positions of the heptad repeat) were leucines to limit their potential contributions to bilin stabilization by structured interactions within the core.

**Figure 2.**
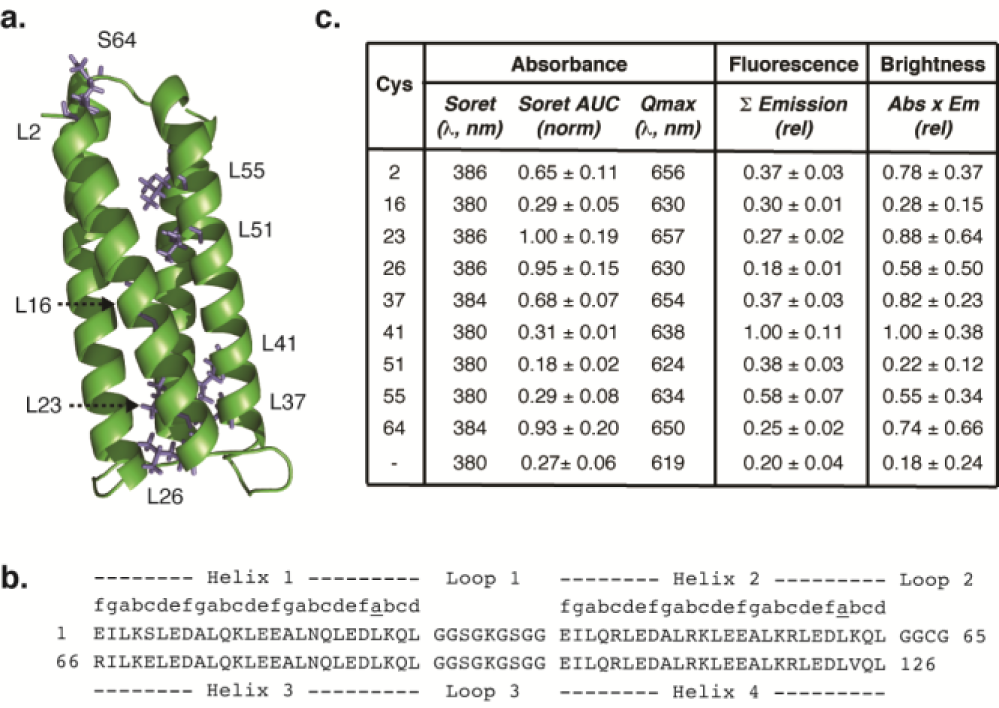
Cysteine scanning for biliverdin (BV) attachment. **(a)** Rosetta-generated Pymol model of the maquette scaffold, with candidate cofactor ligation position **(b)** Sequence of the scaffold, with all hydrophobic core residues (a-d positions) as leucine. **(c)** Summary of relative BV attachment, fluorescence, and brightness of cysteine scanning library (mean ± s.e.). BV attachment quantified from the Soret absorbance band. Fluorescence measured at fixed holoprotein concentrations estimated by BV absorbance (λ_ex_ = 600 nm). Brightness calculated as absorbance *x* fluorescence. (AUC = area under curve, Abs = absorbance, Em = emission, Q = Q-band).

BV attachment trended with cysteine solvent exposure, where those in the solvent exposed B-loop (S64C) or near the termini (L23C) provided good relative balances of appreciable cofactor attachment and baseline fluorescence without stabilization beyond partitioning into the hydrophobic core (**Figure 2c**). In selecting initial starting points for further molecular engineering, we prioritized BV attachment efficiency over initial fluorescence given the reported challenges in cofactor uptake into Bph-FPs^17, 19, 23^. Subsequent engineering proceeded more quickly with the S64C maquette, and thus proteins reported hereon come from the loop-bound starting point. Rosetta modeling of the holoprotein suggested a favored BV placement in the core such that an existing arginine (R119) and lysine (K77) of the scaffold stabilize the BV propionates (similar to Figure 3a-b).

**Figure 3.**
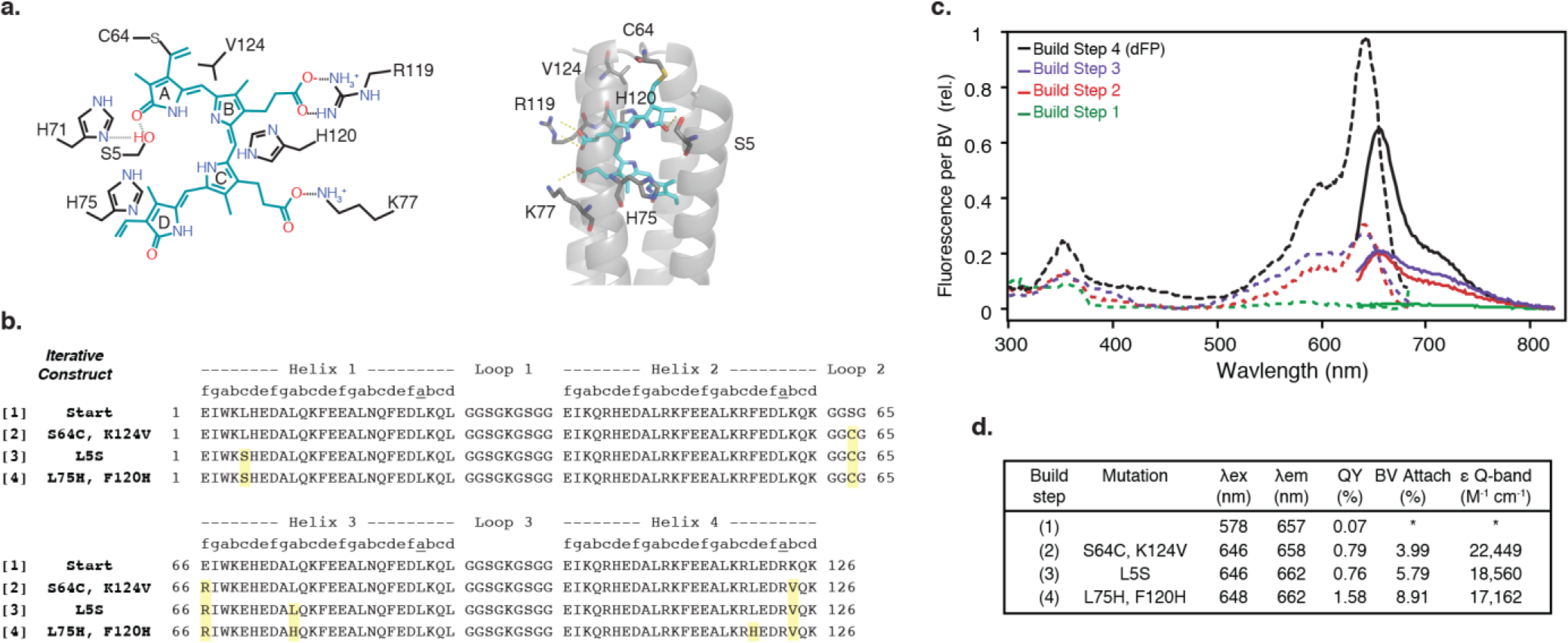
Rational engineering of a biliverdin-binding *de novo*-designed fluorescence-activating protein (dFP). **(a)** (left) Homology-based contact schematic for biliverdin (BV) stabilization within the core (black = side-chains, green = BV), and (right) Pymol visualization of BV binding site in Rosetta-modeled protein core, showing potential stabilization contacts. **(b)** Sequence alignment of the build series, with mutated residues in yellow. The E66R mutation was introduced with S64C based on “CXR” motifs of natural bili-proteins, but did not contribute to stabilization. **(c)** Excitation (dashed line, λem > 715 nm) and emission spectra (solid line, λex = 600 nm) of the step-wise construction, resulting in dFP. **(d)** Photophysical summary of the build series. QY = relative quantum yield vs. Cy5. ε = extinction coefficient. % = below quantification limit. (b-d) Build step 4 = dFP.

### Rational cofactor stabilization

**A** series of stepwise modifications (**Figure 3a-b**) had the intended hierarchical effects of increasing BV attachment efficiency, enhancing fluorescence quantum yield (ϕ_F_), and sharpening the Q-band peak of the bilin absorbance spectra (**Figure 3c-d** and **Supplementary Figure 1**), with the latter two events indicative of bilin rigidification^40^. The helix 4 terminus adjacent to the BV-binding cysteine (C64) was rigidified and made more hydrophobic by strategically positioning a valine (K124V) at the interfacial b-position of the last heptad repeat (Build Step 2 in Figure 3). BV stabilization and placement continued from the A-ring, as the most defined pyrrole in spatial position due to its covalent cysteine attachment, by introducing a serine residue (L5S) intended to hydrogen bond the A-ring nitrogen (Build Step 3).

The cofactor B−, C−, and D-rings were further immobilized by positioning histidine residues (L75H and F120H) to pi-stack to the pyrrole rings and provide hydrophobic core bulk to restrict protein movement and core water access (Build Step 4). Such introduction of aromatic core bulk has previously stabilized similar maquettes that bind tetrapyrroles, as evident by increased melting temperatures and reduced helical movement^7, 9^. Rosetta modeling of the final product suggests that S5 may hydrogen bond to the oxygen of the A-ring, and that H71 also hydrogen bonds to S5 to further stabilize the A-ring through increased steric constraint (**Figure 3a-b**).

The resultant 15-kD monomer fluoresced modestly in the far-red spectrum (λ_ex_ = 648 nm, λ_em_ = 662 nm, ϕ_F_ = 1.58%). This quantum yield is similar to sandercyanin, a naturally occurring fluorescent BV-binding fish pigment (ϕ_F_ = 1.6%)^41^, and less than those of BV-binding directed evolution products derived from Bph‐ and AP (ϕ_F_ ~ 7-18%)^17-23, 31^. For simplicity, we hereon call this *de novo*-designed fluorescence-activating protein, “dFP.”

The holoprotein was predominantly monomeric in analytical ultracentrifugation (AUC) assays (**Supplementary Figure 1**). Circular dichroism (CD) measurements confirmed the helicity of the bundle, and showed that cofactor binding enhanced the overall thermal stability of the protein (T_m-apo_ = 44.7°C *vs*. T_m-holo_ = 50.8°C) (**Supplementary Figure 2**). Electrospray ionization mass spectrometry (ESI/MS) and zinc acetate-staining of denaturing protein gels confirmed covalent attachment of BV to the scaffold (**Supplementary Figure 3a-c**), and filtration on desalting columns was sufficient to remove excess BV adsorbed to the scaffold exterior (**Supplementary Figure 3d**).

As intended, biliverdin covalently attached to the cysteine of the protein scaffold by thioether formation at the vinyl position of the cofactor based on acidic denaturation studies (**Supplementary Figure 4a-d**). The dFP(C64S) mutation destabilized the BV within the core, based on diminished Q-band absorbance and quantum yield (ϕ_F_ = 0.8%) (**Supplementary Figure 4b**). The mutation also reduced the uptake of BV cofactor, which was non-covalently core-bound because it was lost upon acidic denaturation in guanidinium chloride and column filtration. Similarly, the cysteine-containing dFP bound less mesobiliverdin IXa (mesoBV) cofactor, which differs from BV in the reduction of the vinyl sidechains to ethyl groups, and acidic denaturation and column filtration of the mesoBV-bound holoprotein stripped the cofactor from the protein scaffold (**Supplementary Figure 4c**).

The covalently attached BV of denatured dFP did not appreciably photoconvert upon stimulation (λ = 610 +/−5 nm or > 650 nm) based on the difference spectrum (**Supplementary Figure 5**). This result suggests that the cofactor D-ring adopts a 15*Z* configuration^42^ as designed (schematized in **Figures 1a and 3b**) and modeled in Rosetta, and not the photoconverting 15*E* configuration^42^. A future high-resolution structure would greatly inform the structure-function of dFP and related holoproteins.

### Miniature 10 kD bili-protein (“mini”)

While the *de novo*-designed bili-protein is already compact, the 15-kD scaffold contains extraneous heptad repeats, with respect to the needs for cofactor binding and stabilization (**Figure 4a**). These repeats can theoretically be removed without major risk of misfolding in a maquette, as a self-assembling bundle with fairly independent structure-function between the individual residues. Thus, we sought to further push the limit of compactness for a single-chain bili-protein, by engineering a miniature dFP (here called “dFP-mini” or “mini”) by truncating the heptad repeats down to where homology modeling predicts all four pyrrole rings of BV are solvent shielded, while still allowing for the bundle to self-assemble stably (**Figure 4a**). To preserve bundle stability, truncation began after the hydrophobic caps at the helical termini nearest to the D-ring, and Loops 1 and 3 were also shortened. A 10-kD mini, whose helices extended a half-heptad repeat beyond the furthest modeled D-ring contact residue, was stable whereas shorter proteins that terminated at the final hypothetical D-ring contact were unstable.

**Figure 4.**
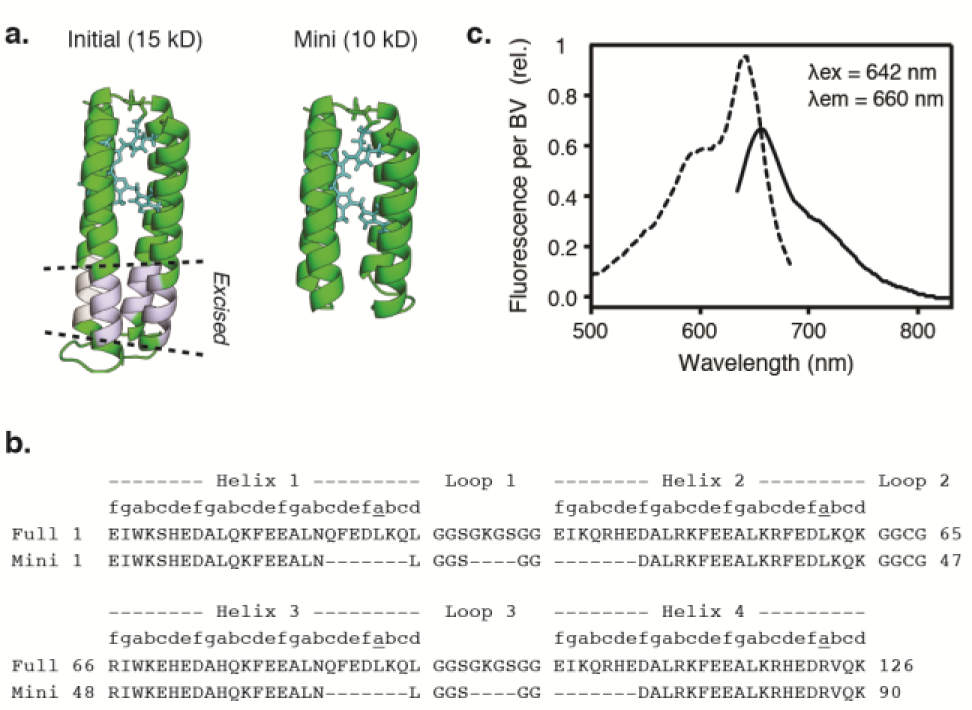
A miniature 10 kD *de novo*-designed bili-protein (mini). **(a)** Pymol render of the BV binding site in Rosetta-modeled full-length and mini bili-protein. **(b)** Sequence alignment, with helix position a-g denoted. **(c)** Excitation (dashed line, λ_em_ > 715 nm) and emission spectra (solid line, λ_ex_ = 600 nm) of the mini (Quantum yield (ϕ_f_) = 1.48%, α Q-band = 16,209 cm^-1^ M^-1^, 16.44% attachment efficiency).

Consistent with the general modularity found amongst maquettes, the mini showed nearly identical photophysical properties to the full-length dFP (**Figure 4b**, λ_ex_ = 642 nm, λ_em_ = 660 nm, ϕ_F_ = 1.48%). The mini also covalently bound the cofactor by thioether formation at the vinyl position, as determined similarly to the full-length scaffold by mass spectrometry and binding/denaturing studies with a cysteine-to-serine mutant (C46S) and meso-BV (**Supplementary Figure 4**). Likewise, the denatured mini did not photoconvert, suggesting that the cofactor D-ring adopts the intended 15*Z* D-ring orientation (**Supplementary Figure 5**).

For reference, the mini is approximately one-third the size of canonical GFP (27 kD) and monomeric Bph-FPs (30-35 kD)^18, 21^, and half the size of a minimal fluorescent domain engineered from a Bph GAF domain (18 kD)^20^. The facile truncation and overall reduction in size compared to Bph-derived fluorochromes reflects the structural simplicity of the *de novo*-designed scaffold. Beyond the three inter-helical loops and hydrophobic caps, as a single protein fold, the mini essentially lacks accessory structural elements other than the cofactor-binding pocket itself.

Compact protein fluorochromes are advantageous as components of molecular sensors, because the size facilitates shortening Förster distances in fluorescence resonance energy transfer (FRET)^43^ and limits potential interference with the activity of fusion partners. Given that BV is endogenously found in eukaryotes, this study is a valuable step toward the potential creation of fully genetically encoded *de novo*-designed reporters, with primary next steps of increased quantum yield and bilin uptake to enable robust performance.

The uptake reported here reflects an *in vitro* autocatalytic attachment efficiency, in the absence of an evolutionarily conserved bilin lyase domain (BLD)^38-40^ or accessory bilin lyase to promote attachment^44-46^. To the best of our knowledge, the analogous efficiencies of engineered Bph-derived fluorescent proteins (i.e. before chromatographic separation of apoprotein from holoprotein) have not been reported. The attachment process to the *de novo*-designed scaffold likely begins with cofactor partitioning into the core and stabilization within the binding pocket preceding thioether formation as described for natural phytochromes^38, 47^ – given that the cysteine-to-serine mutations reduced but did not eradicate holoprotein formation, and that attachment efficiency trended with quantum yield.

Thus, we anticipate that improved cofactor stabilization or rigidification within the core will promote thioether formation and consequently enhance attachment efficiency, fluorochrome absorbance, and quantum yield. These enhancements may arise from complementary approaches in directed evolution and *de novo* design, including the use of new design tools, such as a very recently reported rotamer interaction field (RIF) algorithm that permits the decoupling of ligand-docking optimization in Rosetta from the optimization of the overall scaffold backbone^27^. RIF-guided design begot a *de novo*-designed beta-barrel fluorescence-activating protein^27^ that binds exogenously supplied DFHBI (the chromophore of canonical green fluorescent protein), with a modest quantum yield (ϕ_F_ = 2%) similar to those for the *de novo*-designed biliproteins here.

To summarize, through a rational and step-wise build aided by energy minimization modeling, we constructed highly compact *de novo* proteins that covalently bound and stabilized BV. In keeping with the tenets of synthetic biology and protein design, these self-assembling holoproteins were built from the bottom-up from first principles rather than engineered from the top-down using natural protein starting points.

## ASSOCIATED CONTENT

### Materials and Methods

Refer to Supporting Information for experimental procedures. Plasmids will be made available at Addgene.

### Supporting Information

Materials and Methods. Supplementary Figure 1: Cofactor rigidification and scaffold stabilization by holoprotein formation. Supplementary Figure 2. Covalent attachment of BV to monomeric bundles.

## AUTHOR INFORMATION

### Corresponding Author

* Address: 210 South 33^rd^ Street, Skirkanich Hall Suite 240, Philadelphia, PA 19104, United States. Phone: (215) 898-5159.

### Author Contributions

All authors contributed to experiment and protein design, data analysis, and manuscript preparation. MMS, MSM, IAK, JAM, GK, and CCM conducted protein characterization. IAK and MSM built the Rosetta models. CCM, PLD, and BYC coordinated research.

### Funding Sources

BYC was supported by: National Institutes of Health (1R21DA040434, 1R21EY027562, 1R01NS101106), National Science Foundation (CBET 126497, CAREER MCB 1652003), and the University of Pennsylvania’s University Research Foundation. PLD was supported by Department of Energy (DESC0001035). MSM was supported by the NSF GRFP Fellowship. IAK was supported by the Paul and Daisy Soros Fellowship for New Americans.

## ACKNOWLEDGMENTS

We thank Kushol Gupta (analytical ultracentrifugation), and Leland Mayne and Kendrick Laboratories (mass spectrometry) for technical assistance. We also thank Donald Bryant, J. Clark Lagarias, Katrina Forest, Nathan Rockwell, and Vladislav Verkhusha for helpful discussion on bili-protein structure-function and analysis methods.

## REFERENCES

1. Huang, P.-S.; Boyken, S. E.; Baker, D., The coming of age of de novo protein design. Nature 2016, 537 (7620), 320-327.

2. Bryson, J. W.; Betz, S. F.; Lu, H. S.; Suich, D. J.; Zhou, H. X.; O'Neil, K. T.; DeGrado, W. F., Protein design: a hierarchic approach. Science 1995, 270 (5238), 935-41.

3. Dahiyat, B. I.; Mayo, S. L., De novo protein design: fully automated sequence selection. Science 1997, 278 (5335), 82-7.

4. Beasley, J. R.; Hecht, M. H., Protein design: the choice of de novo sequences. The Journal of biological chemistry 1997, 272 (4), 2031-4.

5. Regan, L.; DeGrado, W., Characterization of a helical protein designed from first principles. Science 1988, 241 (4868), 976-978.

6. Robertson, D. E.; Farid, R. S.; Moser, C. C.; Urbauer, J. L.; Mulholland, S. E.; Pidikiti, R.; Lear, J. D.; Wand, A. J.; DeGrado, W. F.; Dutton, P. L., Design and synthesis of multi-haem proteins. Nature 1994, 368 (6470), 425-32.

7. Farid, T. A.; Kodali, G.; Solomon, L. A.; Lichtenstein, B. R.; Sheehan, M. M.; Fry, B. A.; Bialas, C.; Ennist, N. M.; Siedlecki, J. A.; Zhao, Z. Y.; Stetz, M. A.; Valentine, K. G.; Anderson, J. L. R.; Wand, A. J.; Discher, B. M.; Moser, C. C.; Dutton, P. L., Elementary tetrahelical protein design for diverse oxidoreductase functions. Nature Chem. Biol. 2013, 9 (12), 826-833.

8. Mancini, J. A.; Kodali, G.; Jiang, J.; Reddy, K. R.; Lindsey, J. S.; Bryant, D. A.; Dutton, P. L.; Moser, C. C., Multi-step excitation energy transfer engineered in genetic fusions of natural and synthetic light-harvesting proteins. Journal of The Royal Society Interface 2017, 14 (127).

9. Mancini, J. A.; Sheehan, M.; Kodali, G.; Chow, B. Y.; Bryant, D. A.; Dutton, P. L.; Moser, C. C., De novo synthetic biliprotein design, assembly and excitation energy transfer. Journal of The Royal Society Interface 2018, 15 (141).

10. Sharp, R. E.; Moser, C. C.; Rabanal, F.; Dutton, P. L., Design, synthesis, and characterization of a photoactivatable flavocytochrome molecular maquette. Proceedings of the National Academy of Sciences 1998, 95 (18), 10465.

11. Hecht, M. H.; Richardson, J. S.; Richardson, D. C.; Ogden, R. C., De novo design, expression, and characterization of Felix: a four-helix bundle protein of native-like sequence. Science 1990, 249 (4971), 884.

12. Digianantonio, K. M.; Hecht, M. H., A protein constructed de novo enables cell growth by altering gene regulation. Proc. Natl. Acad. Sci. U.S.A. 2016, 113 (9), 2400-2405.

13. Huang, P. S.; Oberdorfer, G.; Xu, C.; Pei, X. Y.; Nannenga, B. L.; Rogers, J. M.; DiMaio, F.; Gonen, T.; Luisi, B.; Baker, D., High thermodynamic stability of parametrically designed helical bundles. Science 2014, 346 (6208), 481-5.

14. Murphy, G. S.; Sathyamoorthy, B.; Der, B. S.; Machius, M. C.; Pulavarti, S. V.; Szyperski, T.; Kuhlman, B., Computational de novo design of a four-helix bundle protein—DND_4HB. Protein Science 2014, 24 (4), 434-445.

15. Polizzi, N. F.; Wu, Y.; Lemmin, T.; Maxwell, A. M.; Zhang, S.-Q.; Rawson, J.; Beratan, D. N.; Therien, M. J.; DeGrado, W. F., Strategy for designing hyperstable, non-natural protein-ligand complexes with sub-Å accuracy. Nature chemistry 2017, 9 (12), 1157-1164.

16. O'Neil, K. T.; DeGrado, W. F., A thermodynamic scale for the helix-forming tendencies of the commonly occurring amino acids. Science 1990, 250 (4981), 646-51.

17. Shu, X.; Royant, A.; Lin, M. Z.; Aguilera, T. A.; Lev-Ram, V.; Steinbach, P. A.; Tsien, R. Y., Mammalian expression of infrared fluorescent proteins engineered from a bacterial phytochrome. Science 2009, 324 (5928), 804-7.

18. Yu, D.; Baird, M. A.; Allen, J. R.; Howe, E. S.; Klassen, M. P.; Reade, A.; Makhijani, K.; Song, Y.; Liu, S.; Murthy, Z.; Zhang, S. Q.; Weiner, O. D.; Kornberg, T. B.; Jan, Y. N.; Davidson, M. W.; Shu, X., A naturally monomeric infrared fluorescent protein for protein labeling in vivo. Nature methods 2015, 12 (8), 763-5.

19. Filonov, G. S.; Piatkevich, K. D.; Ting, L. M.; Zhang, J.; Kim, K.; Verkhusha, V. V., Bright and stable near-infrared fluorescent protein for in vivo imaging. Nature biotechnology 2011, 29 (8), 757-61.

20. Rumyantsev, K. A.; Shcherbakova, D. M.; Zakharova, N. I.; Emelyanov, A. V.; Turoverov, K. K.; Verkhusha, V. V., Minimal domain of bacterial phytochrome required for chromophore binding and fluorescence. Scientific Reports 2015, 5, 18348.

21. Shcherbakova, D. M.; Baloban, M.; Emelyanov, A. V.; Brenowitz, M.; Guo, P.; Verkhusha, V. V., Bright monomeric near-infrared fluorescent proteins as tags and biosensors for multiscale imaging. Nature communications 2016, 7.

22. Shcherbakova, D. M.; Baloban, M.; Verkhusha, V. V., Near-infrared fluorescent proteins engineered from bacterial phytochromes. Curr Opin Chem Biol 2015, 27, 52-63.

23. Bhattacharya, S.; Auldridge, M. E.; Lehtivuori, H.; Ihalainen, J. A.; Forest, K. T., Origins of Fluorescence in Evolved Bacteriophytochromes. Journal of Biological Chemistry 2014.

24. Kaberniuk, A. A.; Shemetov, A. A.; Verkhusha, V. V., A bacterial phytochrome-based optogenetic system controllable with near-infrared light. Nature methods 2016, 13, 591.

25. Meiler, J.; Baker, D., ROSETTALIGAND: protein-small molecule docking with full side-chain flexibility. Proteins 2006, 65 (3), 538-48.

26. Rohl, C. A.; Strauss, C. E.; Misura, K. M.; Baker, D., Protein structure prediction using Rosetta. Methods Enzymol 2004, 383, 66-93.

27. Dou, J.; Vorobieva, A. A.; Sheffler, W.; Doyle, L. A.; Park, H.; Bick, M. J.; Mao, B.; Foight, G. W.; Lee, M. Y.; Gagnon, L. A.; Carter, L.; Sankaran, B.; Ovchinnikov, S.; Marcos, E.; Huang, P.-S.; Vaughan, J. C.; Stoddard, B. L.; Baker, D., De novo design of a fluorescence-activating β-barrel. Nature 2018.

28. Lehtivuori, H.; Bhattacharya, S.; Angenent-Mari, N. M.; Satyshur, K. A.; Forest, K. T., Removal of Chromophore-Proximal Polar Atoms Decreases Water Content and Increases Fluorescence in a Near Infrared Phytofluor. Frontiers in molecular biosciences 2015, 2, 65.

29. Murphy, J. T.; Lagarias, J. C., The phytofluors: a new class of fluorescent protein probes. Current Biology 1997, 7 (11), 870-876.

30. Gryczynski, I.; Piszczek, G.; Lakowicz, J. R.; Lagarias, J. C., Two-photon excitation of a phytofluor protein. Journal of Photochemistry and Photobiology A: Chemistry 2002, 150 (1–3), 13-19.

31. Rodriguez, E. A.; Tran, G. N.; Gross, L. A.; Crisp, J. L.; Shu, X.; Lin, J. Y.; Tsien, R. Y., A farred fluorescent protein evolved from a cyanobacterial phycobiliprotein. Nat Meth 2016, 9, 763-9.

32. Glazer, A. N.; Stryer, L., Phycofluor probes. Trends in biochemical sciences 1984, 9 (10), 423-427.

33. Grabowski, J.; Gantt, E., Photophysical Properties of Phycobiliproteins from Phycobilisomes: Fluorescence Lifetimes, Quantum Yields, and Polarization Spectra. Photochemistry and photobiology 1978, 28 (1), 39-45.

34. Kumagai, A.; Ando, R.; Miyatake, H.; Greimel, P.; Kobayashi, T.; Hirabayashi, Y.; Shimogori, T.; Miyawaki, A., A bilirubin-inducible fluorescent protein from eel muscle. Cell 2013, 153 (7), 1602-11.

35. Gruber, D. F.; Gaffney, J. P.; Mehr, S.; DeSalle, R.; Sparks, J. S.; Platisa, J.; Pieribone, V. A., Adaptive Evolution of Eel Fluorescent Proteins from Fatty Acid Binding Proteins Produces Bright Fluorescence in the Marine Environment. PLOS ONE 2015, 10 (11), e0140972.

36. Huang, S. S.; Koder, R. L.; Lewis, M.; Wand, A. J.; Dutton, P. L., The HP-1 maquette: From an apoprotein structure to a structured hemoprotein designed to promote redox-coupled proton exchange. Proceedings of the National Academy of Sciences of the United States of America 2004, 101 (15), 5536.

37. Lichtenstein, B. R.; Farid, T. A.; Kodali, G.; Solomon, L. A.; Anderson, J. L. R.; Sheehan, M. M.; Ennist, N. M.; Fry, B. A.; Chobot, S. E.; Bialas, C.; Mancini, J. A.; Armstrong, C. T.; Zhao, Z. Y.; Esipova, T. V.; Snell, D.; Vinogradov, S. A.; Discher, B. M.; Moser, C. C.; Dutton, P. L., Engineering oxidoreductases: maquette proteins designed from scratch. Biochemical Society Transactions 2012, 40, 561-566.

38. Lamparter, T.; Carrascal, M.; Michael, N.; Martinez, E.; Rottwinkel, G.; Abian, J., The Biliverdin Chromophore Binds Covalently to a Conserved Cysteine Residue in the N-Terminus of Agrobacterium Phytochrome Agp1. Biochemistry 2004, 43 (12), 3659-3669.

39. Lamparter, T.; Michael, N.; Caspani, O.; Miyata, T.; Shirai, K.; Inomata, K., Biliverdin Binds Covalently to Agrobacterium Phytochrome Agp1 via Its Ring A Vinyl Side Chain. Journal of Biological Chemistry 2003, 278 (36), 33786-33792.

40. Wu, S. H.; Lagarias, J. C., Defining the bilin lyase domain: lessons from the extended phytochrome superfamily. Biochemistry 2000, 39 (44), 13487-95.

41. Ghosh, S.; Yu, C.-L.; Ferraro, D. J.; Sudha, S.; Pal, S. K.; Schaefer, W. F.; Gibson, D. T.; Ramaswamy, S., Blue protein with red fluorescence. Proceedings of the National Academy of Sciences 2016, 113 (41), 11513.

42. Shang, L.; Rockwell, N. C.; Martin, S. S.; Lagarias, J. C., Biliverdin amides reveal roles for propionate side chains in bilin reductase recognition and in holophytochrome assembly and photoconversion. Biochemistry 2010, 49 (29), 6070-6082.

43. Luedtke, N. W.; Dexter, R. J.; Fried, D. B.; Schepartz, A., Surveying polypeptide and protein domain conformation and association with FlAsH and ReAsH. Nat Chem Biol 2007, 3 (12), 779-784.

44. Biswas, A.; Boutaghou, M. N.; Alvey, R. M.; Kronfel, C. M.; Cole, R. B.; Bryant, D. A.; Schluchter, W. M., Characterization of the Activities of the CpeY, CpeZ, and CpeS Bilin Lyases in Phycoerythrin Biosynthesis in Fremyella diplosiphon Strain UTEX 481. Journal of Biological Chemistry 2011, 286 (41), 35509-35521.

45. Kronfel, C. M.; Kuzin, A. P.; Forouhar, F.; Biswas, A.; Su, M.; Lew, S.; Seetharaman, J.; Xiao, R.; Everett, J. K.; Ma, L.-C.; Acton, T. B.; Montelione, G. T.; Hunt, J. F.; Paul, C. E. C.; Dragomani, T. M.; Boutaghou, M. N.; Cole, R. B.; Riml, C.; Alvey, R. M.; Bryant, D. A.; Schluchter, W. M., Structural and Biochemical Characterization of the Bilin Lyase CpcS from Thermosynechococcus elongatus. Biochemistry 2013, 52 (48), 8663-8676.

46. Wiethaus, J.; Busch, A. W. U.; Kock, K.; Leichert, L. I.; Herrmann, C.; Frankenberg-Dinkel, N., CpeS Is a Lyase Specific for Attachment of 3Z-PEB to Cys82 of β-phycoerythrin from Prochlorococcus marinus MED4. Journal of Biological Chemistry 2010, 285 (48), 37561-37569.

47. Li, L.; Murphy, J. T.; Lagarias, J. C., Continuous Fluorescence Assay of Phytochrome Assembly in Vitro. Biochemistry 1995, 34 (24), 7923-7930.

